# Sequence-dependent molecular asymmetry and architecture define electric potential profiles of biomolecular condensates

**DOI:** 10.64898/2026.08.03.742525

**Authors:** Fangke Chen, Runchen Xia, Yifan Dai, Xiangze Zeng

**Author notes:** Equal contribution (Fangke Chen, Runchen Xia).

## Abstract

Biomolecular condensates, which regulate diverse cellular processes, exhibit distinct electric potential profiles. This potential gradient between the dilute and the dense phases serves as the underlying driving force mediating the unique microenvironment and electrochemical activity of condensates. However, the molecular principles encoding the electric potential profiles of condensates remain unclear. In this study, we show that molecular asymmetry is a unifying origin of electric polarization in condensates. Asymmetric protein–cation and protein–anion affinities alone generate an interfacial electric double layer and a finite potential even in condensates formed by charge-free proteins. The sign of potential gradient follows the direction of the affinity bias, and the magnitude collapses onto a single linear function of dense-phase protein volume fraction across changes in chain length, interaction strength and salt concentration. Further, chain termini preferentially occupy the condensate interface, so charges positioned asymmetrically with respect to the termini create spatial charge separation even in neutral polyampholytes. These interaction-encoded and sequence architecture-encoded asymmetries can reinforce, screen or reverse one another, allowing the magnitude and polarity of the interphase potential to be tuned through sequence design or solvent environments.

## Introduction

Biomolecular condensates formed by phase separation have emerged as important organizers of intracellular space, regulating critical cellular processes ranging from gene transcription and signal transduction to stress responses ^1–8^. Dysregulation of biomolecular condensates has been implicated in severe human diseases, including cancers and neurodegenerative diseases ^3, 9–11^. The functional roles of biomolecular condensates can be broadly grouped into three categories: 1) concentration buffering, which maintains a constant concentrations within the dilute and dense phases for a system within the two-phase regime ^12–14^; 2) reaction crucibles, which concentrate reactants and provide distinct chemical microenvironments that modulate reaction kinetics^15–17^; 3) organizational hubs, which selectively partition or exclude specific biomolecules ^18–23^. This biomolecule-centered view of condensate function has recently begun to shift.

Recent discoveries have demonstrated that condensates are not merely passive organizational hubs but can also act as electrochemically active entities that shape the intracellular electrochemistry ^24^. This electrochemical activity of condensates is thermodynamically defined by their electric potential profiles established by phase transitions of the biomacromolecules. This grand electric potential between the dilute and the dense phases, which is the Galvani potential ^25^, is defined as

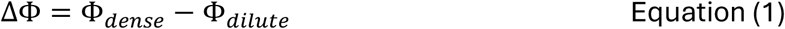

where Φ*_dense_* and Φ*_dilute_* are the electric potentials in the bulk regions of the dense and dilute phases, respectively. This potential is encoded by the molecular compositions within the phase. The potential difference modulates cytoplasmic ion distribution and pH condition ^26, 27^, dictating an interfacial electric field and unique microenvironment. These electrochemical properties of condensates can drive non-enzymatic chemical reactions ^26, 28–30^. Altered condensate electrostatics and interfacial properties have also been implicated in pathological aggregation and may influence toxic self-assembly pathways associated with neurodegenerative diseases^11, 31^.

The establishment of ΔΦ between coexisting phases in equilibrium arises from a complex interplay between different physicochemical properties of condensates, as illustrated in Fig. 1. For a monovalent salt, the relationship among the interphase potential, ion partitioning, and ion-transfer energetics can be expressed through a thermodynamic equation derived from electrochemical equilibrium:

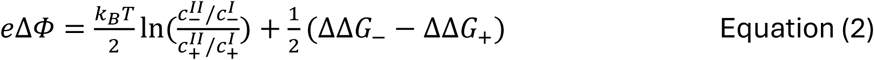

**Figure 1.**
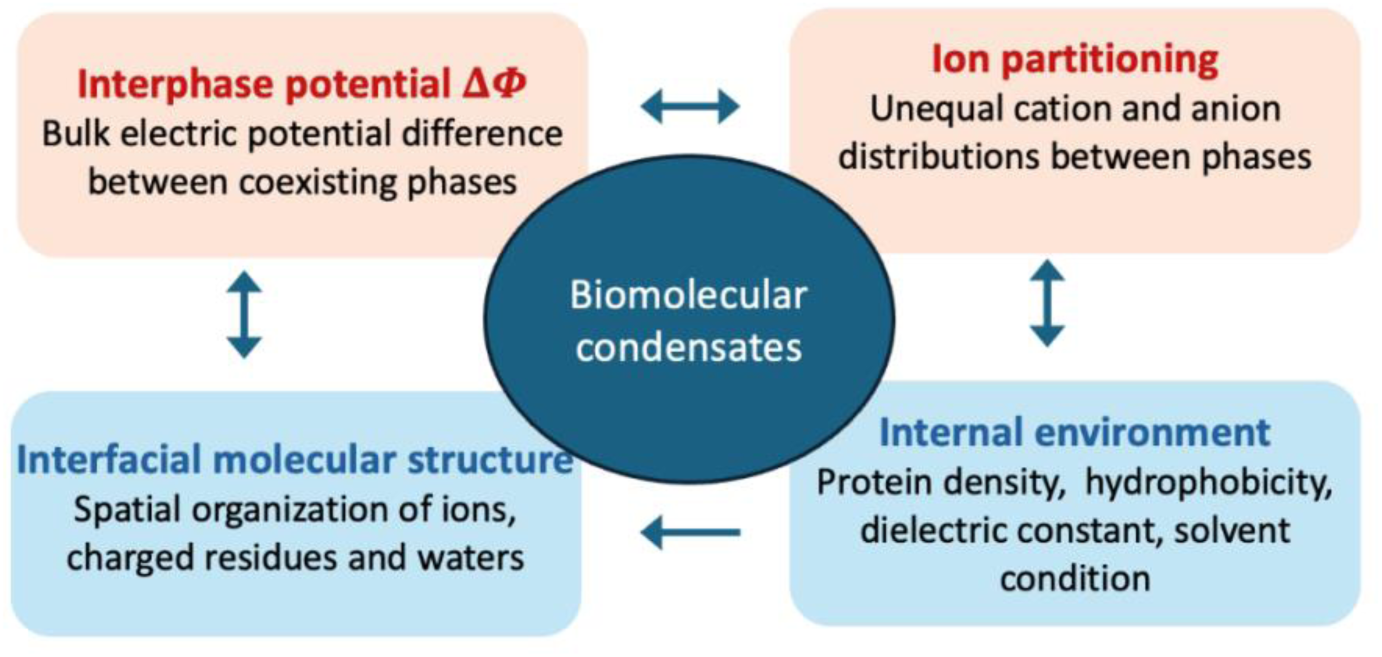
Complex interplay between the physiochemical properties of biomolecular condensates. The interphase electric potential difference, ΔΦ = Φ*_dense_* − Φ*_dilute_*, emerges from the coupled effects of condensate internal environment, ion partitioning, and interfacial molecular organization. Differences in protein concentration and solvent properties affect the transfer energetics and partitioning of cations and anions. Ion partitioning and interfacial organization generate spatial charge and dipole distributions, which determine the electric field and potential profile. In turn, the electrostatic environment can influence ion distributions and molecular orientation, producing coupled feedback among these properties.

where phase *I* and *II* denote the dilute and dense phases, respectively, and 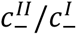 and 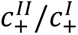 are the partition coefficient of the anion and cation. 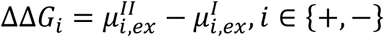 is the intrinsic transfer free energy for ions moving from the dilute to the dense phase. Although Equation 2 provides a macroscopic description, a fundamental gap remains between this thermodynamic framework and its microscopic origins: which molecular interactions and sequence-level features determine ion partitioning, ion-transfer free energies, and ultimately contribute to the magnitude and polarity of the interphase potential in protein condensates? Moreover, beyond the classical Donnan model, where only ions are considered contributors to electric potential ^32^, how do intermolecular and intramolecular interactions among macromolecules within the dense phase also contribute to establishing this potential?

Recent experimental and computational efforts on protein condensates have shown composition-dependent electrochemical properties of condensates. Atomistic simulations revealed asymmetric distribution of salts between the dilute and the dense phases ^33^. Sequence composition of biomolecules was found to modulate the apparent pH of condensates ^26, 34^. The charge profiles of IDPs can encode the capability of condensates to differentially partition cations and anions, thereby establishing distinct interphase potentials ^35, 36^. Further, terminal sequence compositions play a critical role in modulating surface charge density of condensates, thereby affecting interfacial potential gradients ^28, 30^. Interestingly, for condensates formed by charge-free disordered proteins in salt solution, a measurable electric potential was observed^37^. Atomistic simulations and free-energy calculations revealed a pronounced thermodynamic asymmetry between anions and cations (ΔΔ*G*_+_ < 0 < ΔΔ*G*_−_), indicating that transfer of cations into the dense phase is energetically favorable because of preferential protein–cation interactions. Both ΔΔ*G*_+_ and ΔΔ*G*_−_ were found to scale linearly with the dense-phase protein volume fraction (*φ_p_*), identifying dense-phase macromolecular density as a primary driver of the interphase potential.

Related electrochemical phenomena have been investigated theoretically in complex coacervates formed by oppositely charged polyelectrolytes, which is the same formation mechanism as many physiological condensates, such as stress granules ^38^. Zhang et al found that a simple concentration asymmetry between polycation and polyanion can lead to a charge separation at the interface and the interphase potential Δ*Φ* ^25^. They subsequently showed that the interfacial net charge profile and electric potential profile can be tuned by the overall salt concentration, with increasing salt concentration generally reducing the interphase potential ^39^. Similar salt-dependent behavior was reported by Majee et al ^40^, who further showed that the interfacial tension can regulate the net charge profile and the interfacial electric potential profile.

Taken together, these previous works point to molecular asymmetry as a common physical origin of electric potentials of condensates. However, it remains unclear how sequence properties of disordered properties can encode molecular asymmetry, thereby defining distinct electric potential profiles of condensates. An overlooked source of asymmetry is the topological distinction between terminal and interior residues. Compared with interior residues, terminal residues are subject to fewer chain-connectivity constraints and may therefore exhibit an enhanced propensity to localize at a condensate interface. If positive and negative residues are distributed asymmetrically with respect to the two chain termini, this positional preference could produce unequal interfacial enrichment of the two charge types. The resulting charge separation could establish an electric double layer and determine both the sign and magnitude of Δ*Φ*, even when the protein sequence is net neutral. This concept is aligned with previous experimental observations ^28^. However, whether the intrinsic interfacial preference of chain termini is itself sufficient to alter a finite Δ*Φ*, and how this effect depends quantitatively on terminal charge placement, remain unknown.

To address these questions, in this study, we performed systematic coarse-grained molecular dynamics simulations to identify minimal physical principles governing the interphase potential. First, using a minimalist homopolymer model of charge-free proteins in explicit monovalent salt, we show that asymmetric protein-cation and protein-anion interactions are sufficient to generate interfacial charge separation and Δ*Φ*. Across distinct physical perturbations including chain length, protein-protein interaction strength, and salt concentration, the calculated interphase potential collapses onto a common master curve when plotted as a function of protein volume fraction. This master curve exhibits a linear regime consistent with thermodynamic predictions. Next, using neutral disordered proteins under conditions without protein-ion interaction asymmetry, we uncovered a pronounced positional preference of chain termini at the condensate interface. To quantify this effect, we define a sequence-level order parameter Net Charge Preference at chain Terminus (NCPT), which correlates with the magnitude and polarity of Δ*Φ*. Finally, we investigated the competition between sequence-encoded NCPT and protein-ion interaction asymmetry. Our study establishes a quantitative molecular framework for understanding condensate electrochemical properties, illustrating the sequence determinants that can modulate condensate electrochemistry.

## Results

### A minimalist coarse-grained model generates an interphase electric potential in condensates formed by charge-free disordered proteins

We first asked if the interphase potential previously observed in condensates formed by charge-free intrinsically disordered proteins (IDPs) from our previous study ^37^ can be reproduced using a minimalist coarse-grained model. Coarse-grained molecular simulations have become important tools for studying biomolecular phase separation because they access the collective length and time scales of condensate formation while retaining selected molecular or sequence-dependent interactions ^41–48^. Such models have been used to connect chain length, sequence patterning, and residue-level interactions to protein conformations, phase diagrams, and condensate organization ^23,49–58^.

To address this question, we represented each IDP as a homopolymer composed of 20 neutral coarse-grained beads, and included explicit salt ions (Fig. 2A). Short-range nonbonded interactions were described using the standard Lennard-Jones (LJ) potential, whereas electrostatic interactions between ions were modeled by the Columb potential. The protein-protein LJ potential well depth, *ε_p_*, was set as 0.2 kcal/mol, which enabled the protein chains to undergo phase separation at 300 K. The force field parameters for Na^+^ and Cl^-^ ions used in coarse-grained simulations were taken from Lin et al ^59^. and the baseline parameterization contained stronger protein–cation than protein–anion interactions. This interaction asymmetry was consistent with, at a phenomenological level, the preferential protein–cation interactions identified in our previous atomistic simulations ^37^. Additional details of the simulation setup are provided in the Methods section.

**Figure 2.**
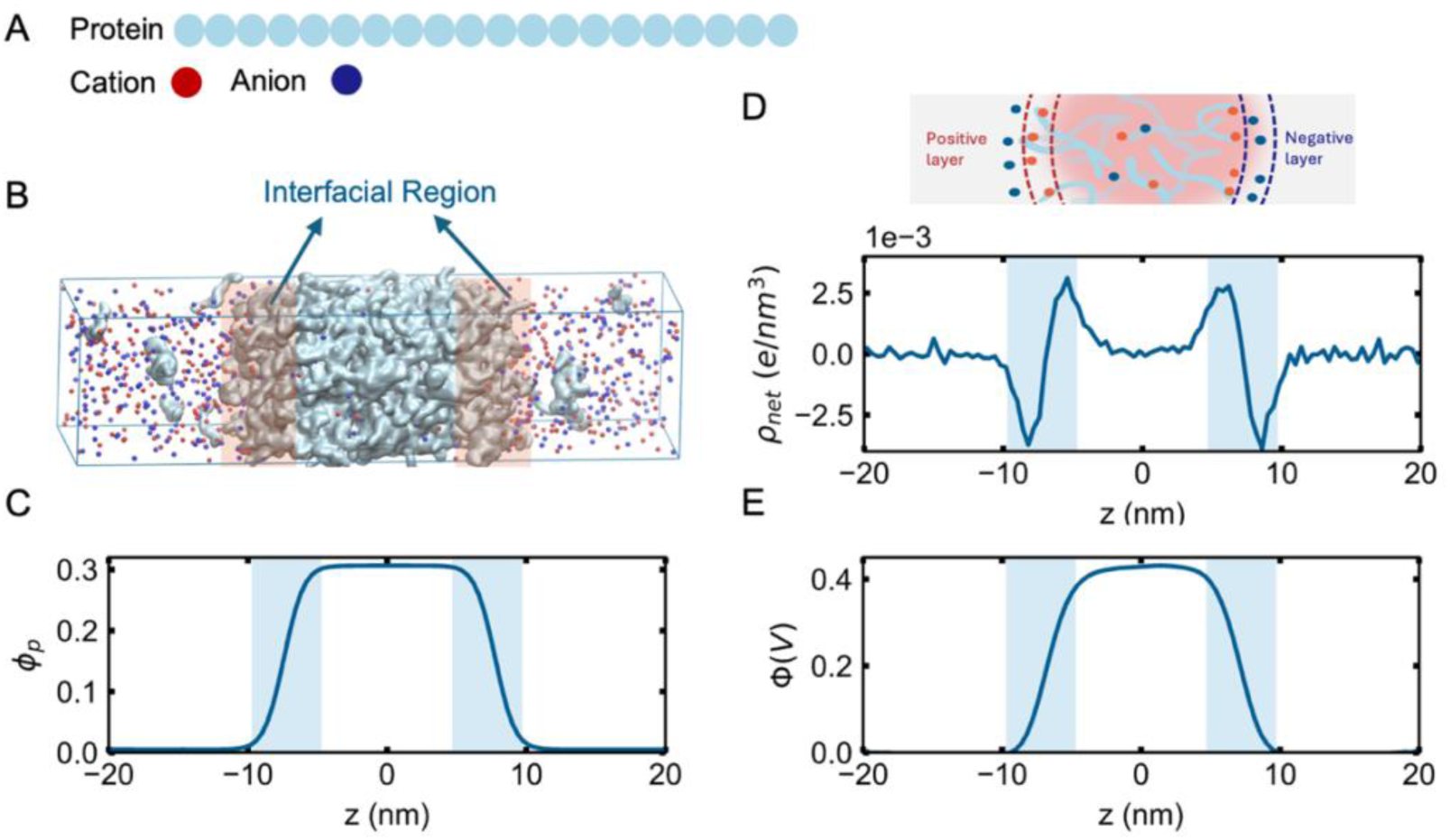
A minimal coarse-grained model generates an interphase electric potential in a charge-free protein condensate. **(A)** Schematic of the coarse-grained model. Each protein is represented as a charge-free homopolymer, whereas monovalent cations and anions are modeled explicitly. Protein– protein and protein–ion short-range interactions are described by Lennard–Jones potentials, and electrostatic interactions between ions are described by the Coulomb potential. **(B)** Representative simulation snapshot showing a protein-rich slab coexisting with a protein-dilute phase. **(C)** Protein volume-fraction profile, *φ_p_*(*z*), along the direction normal to the condensate interface. The central plateau corresponds to the dense phase, and the outer regions correspond to the dilute phase. **(D)** Net charge-density profile, *ρ_net_*(*z*), showing spatially separated positive and negative charge layers at the two condensate interfaces. **(E)** Electric potential profile, Φ(*z*), calculated from the charge-density profile using Poisson’s equation. The difference between the bulk-like dense- and dilute-phase plateaus defines the interphase electric potential, ΔΦ = Φ*_dense_* − Φ*_dilute_*. The dense phase has a higher potential than the dilute phase under the baseline interaction parameters.

Despite its simplicity, this phenomenological model reproduced principal ion-mediated electrochemical features as observed in previous atomistic simulations ^37^, including interfacial charge separation and a finite potential difference between the dense and dilute phases. In slab-geometry simulations, we observed coexisting dilute and dense phases (Fig. 2B), as confirmed by the protein volume fraction along the z direction *φ_p_* (Fig. 2C). We then calculated the net charge density profile along z direction and found a clear charge separation regime localized at the condensate interface, consistent with the formation of an electric double layer (Fig. 2D). Solving Poisson’s equation using the net charge density profile, *ρ_net_*, yielded an electrostatic potential profile Φ(*z*), ^60^ showing that the dense phase is at a higher potential than the dilute phase (Fig. 2E), corresponding to a positive interphase potential ΔΦ = Φ*_dense_* − Φ*_dilute_* > 0. This positive potential profile also matches experimentally measured bulk phase potential gradients using electrochemical potentiometry ^61^. Together, these results demonstrate that a finite ion-mediated interphase potential can emerge in a minimal charge-free polymer model containing asymmetric protein–ion interactions, supporting the use of this minimalist model to study sequence property-dependent condensate potential profiles.

### Asymmetric interactions between cation- and anion-protein generate the interphase potentials

Since the model proteins are electrically neutral, the electric double layer in Fig. 2D cannot arise from fixed charges on the polymer chains themselves. We therefore reasoned that it must originate from asymmetric ion distributions near the condensate surface, which in turn reflect unequal interactions of cations and anions with proteins. To test this idea directly, we systematically varied the cation-protein and anion-protein interaction strengths, denoted by *ε_p_*_+_ and *ε_p_*_−_, respectively, and considered three regimes: *ε_p_*_+_ = *ε_p_*_−_, *ε_p_*_+_ > *ε_p_*_−_, and *ε_p_*_+_ < *ε_p_*_−_ (Fig. 3A).

**Figure 3.**
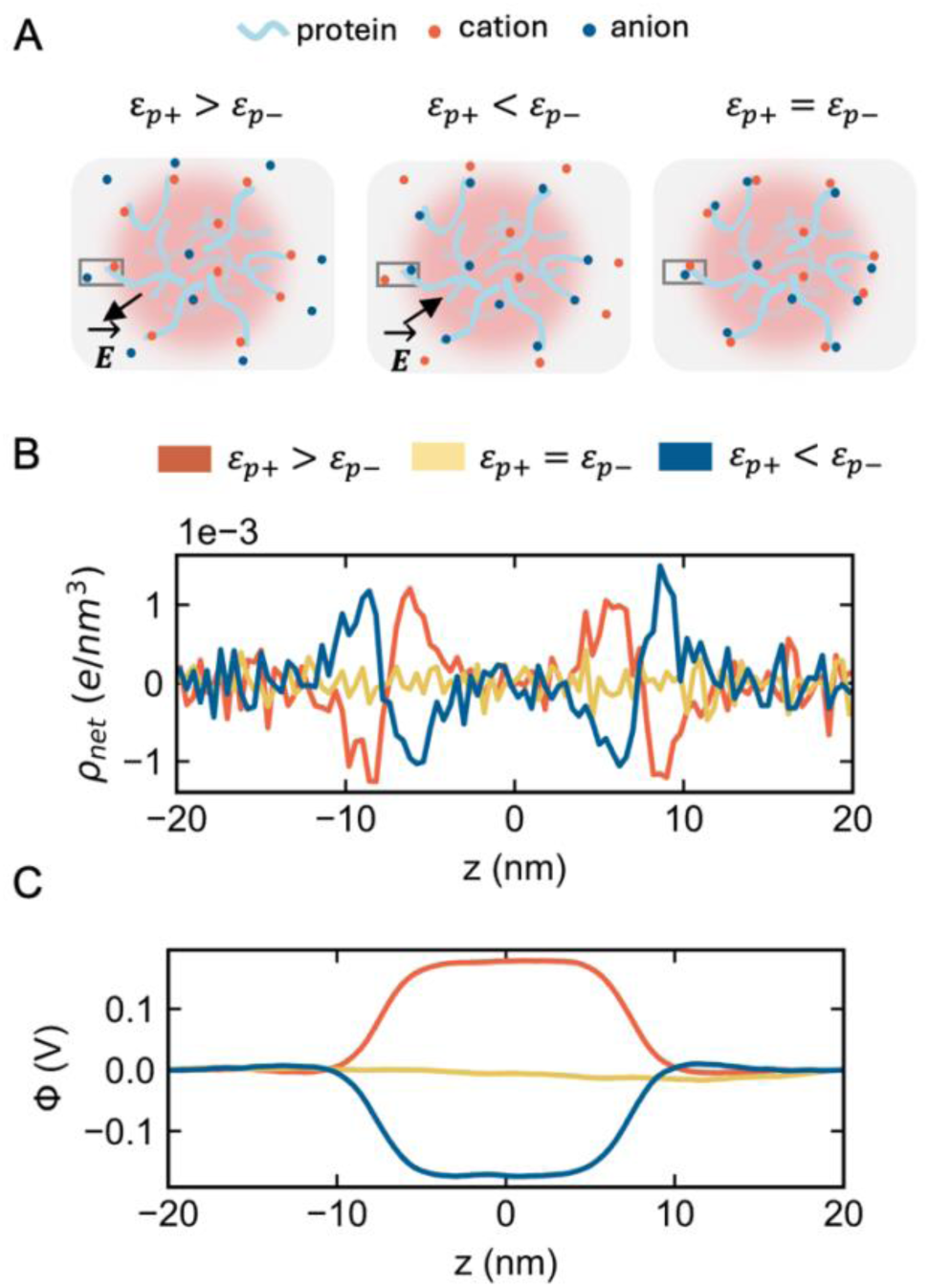
Asymmetric protein–ion interactions determine the formation and direction of the interphase electric potential. **(A)** Schematic of the three protein–ion interaction regimes examined: symmetric interactions, *ε_p_*_+_ = *ε_p_*_−_ ; stronger protein–cation interactions, *ε_p_*_+_ > *ε_p_*_−_ ; and stronger protein–anion interactions, *ε_p_*_+_ < *ε_p_*_−_. **(B)** Net charge-density profiles, *ρ_net_*(*z*), under the three interaction conditions. Symmetric protein–ion interactions produce no persistent interfacial charge separation, whereas asymmetric interactions generate oppositely charged interfacial layers. Reversing the interaction asymmetry reverses the ordering of the charge layers. **(C)** Corresponding electric potential profiles, Φ(*z*). Symmetric interactions yield ΔΦ ≈ 0, whereas unequal protein– cation and protein–anion interactions generate a finite potential. Stronger protein–cation interactions produce ΔΦ > 0, while stronger protein–anion interactions reverse the sign of ΔΦ < 0. These results establish asymmetric protein–ion affinity as a minimal mechanism for interphase-potential generation in charge-free protein condensates.

As expected, when *ε_p_*_+_ = *ε_p_*_−_, the net charge density profile remained essentially flat along the z direction of the simulation box, indicating that no electric double layer forms at the interface (Fig. 3B). Correspondingly, the electric potential in the dense phase is identical to that in the dilute phase (Fig. 3C). In contrast, whenever *ε_p_*_+_ ≠ *ε_p_*_−_, a clear interfacial charge separation emerged accompanied by a finite interphase potential. When *ε_p_*_+_ > *ε_p_*_−_, cations interacted more favorably with the protein-rich phase, producing a positive charge layer on the dense phase-side of the interface and a compensating negative layer on the dilute phase-side. This arrangement yielded a higher electric potential in the dense phase than in the dilute phase. Reversing the interaction asymmetry, such that *ε_p_*_+_ < *ε_p_*_−_, inverted the charge layering and reversed the sign of the interphase potential. These results demonstrate that asymmetric cation- and anion-protein interactions are sufficient to generate the electric double layer and the relative strengths of these interactions determine the direction of the interfacial electric field.

### Dense-phase protein concentration regulates the interphase potential

Because the interfacial charge separation arises from the imbalanced accumulation of cations and anions at the interface, we hypothesized that protein concentration in the dense phase would modulate the extent of ion accumulation and therefore regulate the interphase potential. To test this hypothesis, we systematically tuned the protein volume fraction, which corresponds to the experimentally measured dense-phase protein concentration, using three different approaches: varying protein chain length *N*, increasing protein-protein attraction *ε_pp_*, and changing the overall ion concentration *C_ion_*. All three perturbations altered *φ_p_* and, in turn, the interphase potential (Fig. 4). Increasing the chain length led to a higher protein concentration in the dense phase and a corresponding increase in ΔΦ (Fig. 4A). In contrast, increasing the overall ion concentration reduces both *φ_p_* and ΔΦ (Fig. 4C). This decrease may reflect salt-dependent changes in dense-phase packing and ion partitioning, together with enhanced electrostatic screening of the interfacial charge separation. The dependence on protein-protein interactions was more nuanced: *φ_p_* increased monotonically with *ε_pp_*, whereas ΔΦ first increased and then decreased with a turnover occurring near *φ_p_* ≈ 0.45 (Fig. 4B). A plausible explanation is that, at sufficiently strong protein-protein attraction, ions become increasingly excluded from the dense phase, thereby weakening the interfacial charge separation despite the higher protein concentration.

**Figure 4.**
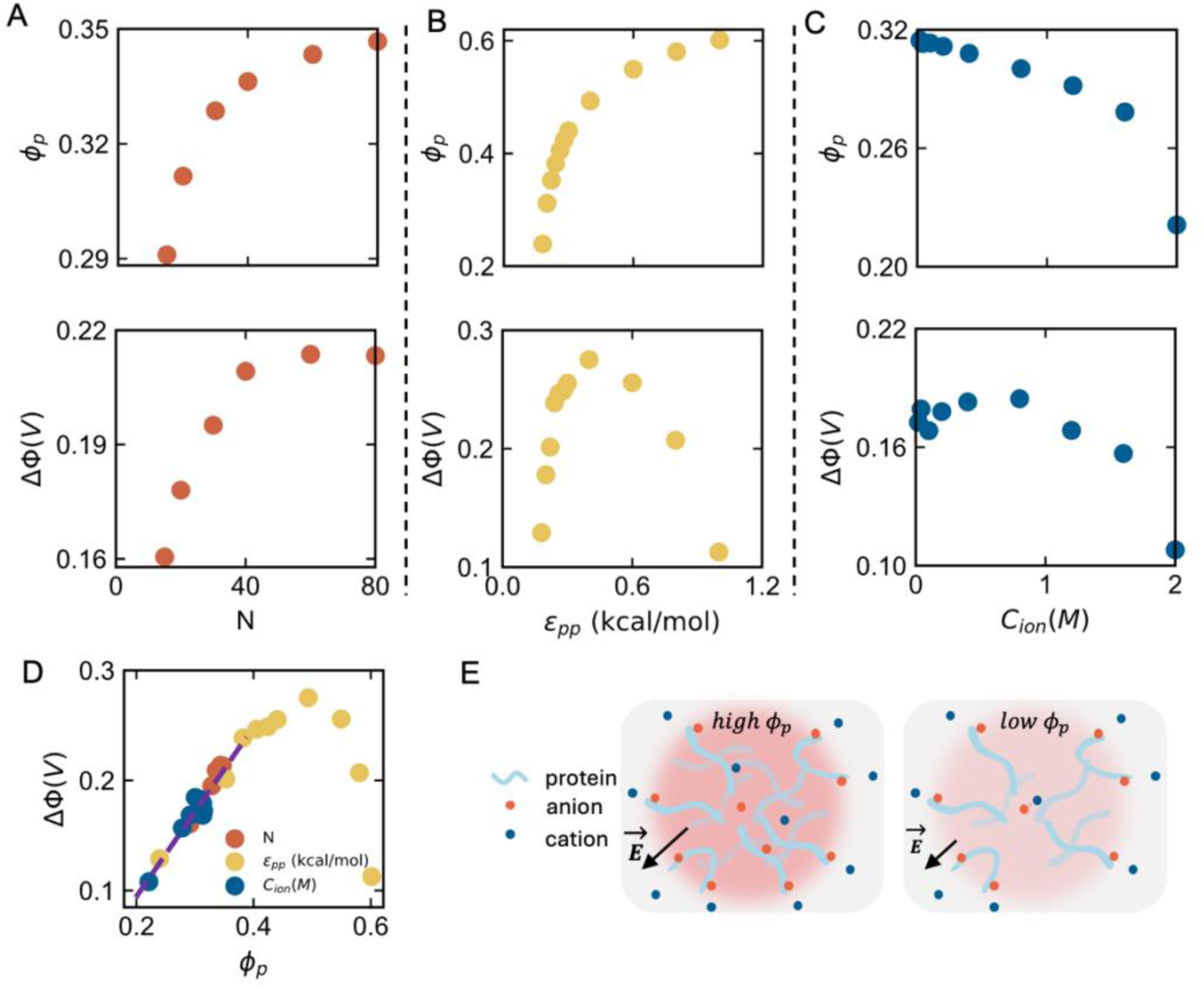
Dense-phase protein volume fraction regulates the interphase electric potential. Dependence of the dense-phase protein volume fraction, *φ_p_*, and interphase electric potential, ΔΦ on protein chain length N **(A),** protein–protein interaction strength *ε_pp_***(B),** and salt concentration *C_ion_* **(C)**. Increasing chain length increases both *φ_p_* and ΔΦ. Increasing protein–protein attraction initially increases ΔΦ, followed by a decrease at high *φ_p_*. Increasing salt concentration reduces both the dense-phase protein volume fraction and the interphase potential under the conditions examined. **(D)** Interphase potential plotted against the dense-phase protein volume fraction for all three perturbations. The data collapse onto a common relationship, indicating that chain length, protein– protein attraction, and salt concentration regulate ΔΦ. At moderate protein volume fractions, ΔΦ depends approximately linearly on *φ_p_*; deviation from linearity occurs at higher volume fractions. **(E)** Schematic illustration of how dense-phase protein volume fraction regulates the interphase potential.

Notably, when ΔΦ was plotted against *φ_p_*, the data obtained from these three approaches collapsed to a common curve (Fig. 4D). This collapse suggests that these distinct control parameters modulate the interphase potential through a common underlying variable, namely the protein concentration within the dense phase. For *φ_p_* < 0.4, ΔΦ increases approximately linearly with *φ_p_*. This linearity can be explained by Eq. 2. For condensates formed by proteins with zero charge, the partitioning term in Eq. 2 is 0 because cation and anion concentrations are equal in the dense phase due to charge neutrality. Thus, the interphase potential is determined entirely by the intrinsic transfer free energies of cations and anions, ΔΔ*G*_−_ and ΔΔ*G*_+_. Our previous free energy calculations showed that both ΔΔ*G*_−_ and ΔΔ*G*_+_ exhibited linear relationships with protein volume fraction (*φ_p_*) in dense phase when *φ_p_* < ∼0.4. Consequently, the interphase potential ΔΦ shows a linear relationship with *φ_p_*. At higher *φ_p_*, the observed deviations from linearity may reflect ion exclusion or a nonlinearity dependence of ion transfer free energies and interfacial organization on protein concentration. Because experimentally measured protein volume fractions in many biomolecular condensates fall below 0.4, this near-linear regime may represent a broadly relevant and experimentally testable feature.

### Terminal charge patterning is a key sequence determinant of interphase potential

The results above focus on condensates formed by uncharged proteins and therefore isolate the contribution of asymmetric ion-protein interactions. We next asked how sequence-encoded charge patterning itself affects the interphase potential. To isolate the contribution of proteins to potential profile, we next simulated the condensate system without salts in the following simulations unless otherwise stated and designed neutral polyampholyte sequences.

Previous studies have established that sequence-dependent intermolecular interactions can regulate condensate interfacial electrostatics ^61^. These studies therefore imply a mechanism in which sequence modifications or changes in interaction strength reorganize the condensate architecture, thereby establishing distinct charge profile across the dilute, the interface, and the dense phases. We first investigated a distinct but complementary mechanism that does not require altering the interaction strengths between terminal and internal protein regions. Specifically, we asked whether the topological preference of chain termini for the condensate interface could translate the positions of charged residues along a sequence into interfacial charge separation.

Indeed, our charge-free homopolymer simulations show that beads near either chain end have a higher probability of localizing at the interface than beads in the chain interior (Fig. S1), which was also observed in previous coarse-grained simulations ^57^. If positive and negative residues occupy nonequivalent positions relative to the two termini, this intrinsic terminal preference can produce unequal interfacial enrichment of the two charge types. The resulting spatial charge separation can generate an interfacial charge double layer and a finite ΔΦ, even when the net charge of the protein sequence is zero.

To examine this mechanism systematically, we designed three sequences with distinct terminal charge arrangements (Fig. 5A). Sequence A1 begins with a positively charged bead and follows an alternating “charged-neutral” pattern along the chain, with the charged sites alternating in sign (+, −, +, −, …), ending with a negatively charged bead. Importantly, the positively charged beads have the mirror symmetry with negatively charged beads with respect to the center of the sequence. Sequence A2 and A3 have one more neutral bead in the C and N terminus of A1, respectively. Thus, the three sequences contain the same periodic arrangement of positive and negative charges, but differ in how those charges are positioned relative to the chain termini.

**Figure 5.**
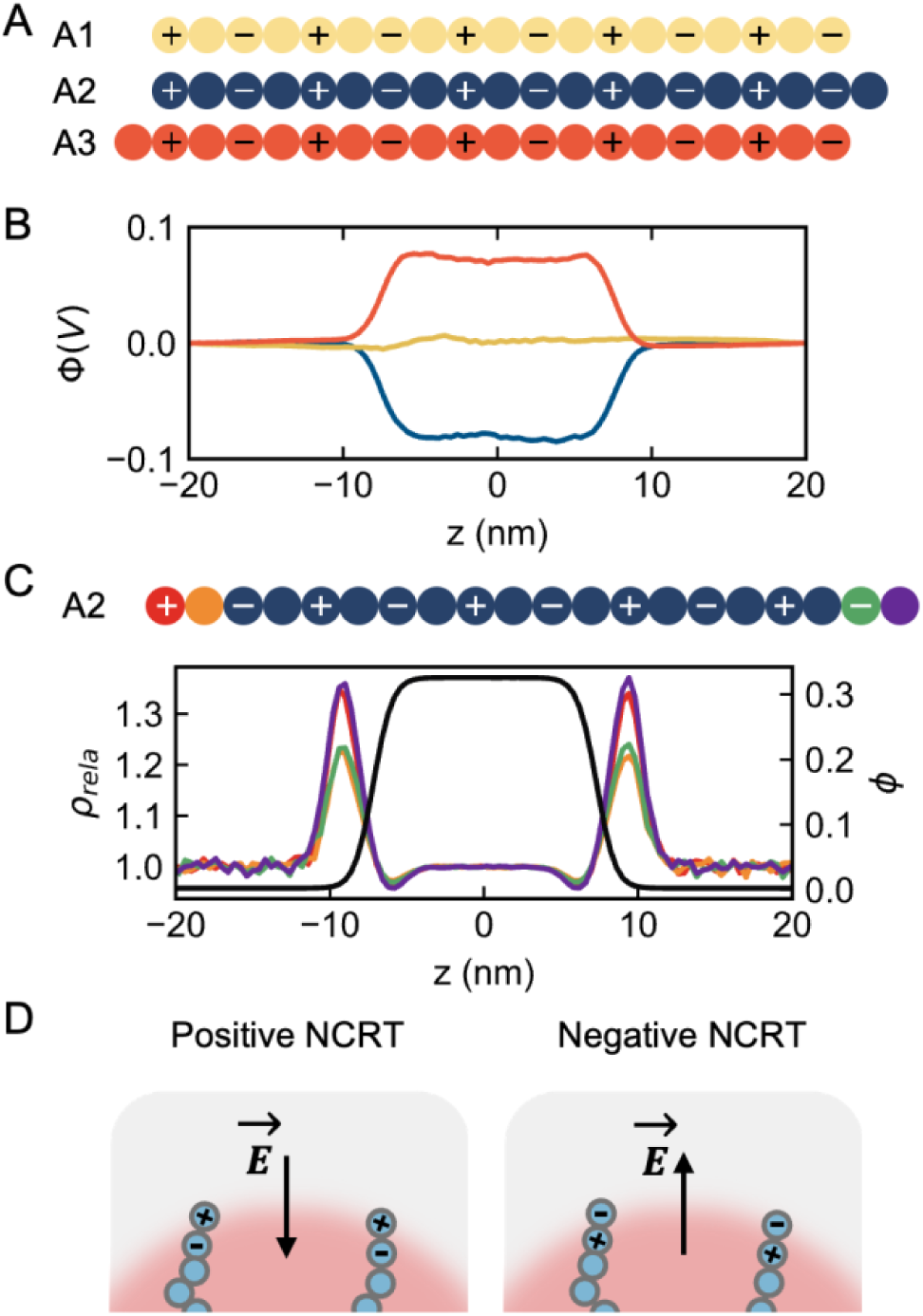
Terminal charge patterning generates interphase electric potentials in neutral polyampholytes. **(A)** Designed net-neutral sequences A1, A2, and A3. The sequences contain related arrangements of positively charged, negatively charged, and neutral beads but differ in the positions of charged residues relative to the chain termini. **(B)** Electric potential profiles for condensates formed by A1, A2, and A3. The terminally symmetric sequence A1 produces negligible interphase potential, whereas A2 and A3 generate finite potentials with opposite signs. **(C)** Relative density profiles of representative terminal and neighboring internal beads in A2. Terminal beads (the N terminal positively charged bead colored by red and C terminal neutral bead colored by purple) exhibit stronger interfacial enrichment than internal beads (the N terminal neutral bead colored by yellow and C terminal negatively charged bead colored by green), producing unequal spatial distributions of positive and negative protein charges. Relative density is defined as the local volume fraction of a selected bead type normalized by the total local protein volume fraction. **(D)** Schematic showing the charge-density profile and corresponding electrostatic organization at the A2 condensate interface. Preferential localization of the positively charged terminal region toward the dilute-phase side generates a positive charge layer on that side and a compensating negative layer toward the dense-phase side. The resulting interfacial charge separation lowers the electric potential of the dense phase relative to the dilute phase. A3 exhibits the opposite terminal charge bias and consequently generates a potential of the opposite sign.

The distinct terminal charge states led to strikingly different interphase potential. For A1, the positive and negative charges are arranged symmetrically with respect to the chain center, thus they display the same propensity for locating at the interface. As a result, we do not observe the electric double layer established at the condensate interface and the interphase potential is essentially the same in the dense and dilute phase (Fig. 5B). In contrast, we observed the electric double layer and interphase potential formed by A2 and A3. While A2 generated a lower electric potential in the dense phase than dilute phase, A3 produced the opposite trend.

To understand the microscopic origin of these behaviors, we analyzed the relative density profiles of the terminal beads for A2 (Fig. 5C). The beads located at the terminus, namely the first positively charged bead and the last neutral bead, showed the greatest enrichment on the dilute-phase side of the interface, whereas the adjacent internal beads, namely the neutral and negatively charged beads showed weaker enrichment. Because the relative density *ρ_rela_* is defined as the volume fraction normalized by the total protein volume fraction along z direction, this imbalance indicates that A2 creates an excess positive charge layer on the dilute-phase side of the interface, and a compensating negative charge layer on the dense-phase side. This charge separation leads to the establishment of the electric double layer with an electric field pointing from the dilute phase to the dense phase (Fig. 5D), thereby making the dense phase electrostatically lower in potential. The opposite sign observed for A3 can be explained by the same mechanism operating with reversed terminal charge bias. These results show that terminal charge patterning alone, even at fixed overall sequence composition and net charge, can dictate both the magnitude and sign of the interphase potential.

### Net Charge Preference at chain Terminus (NCPT) quantitatively describes terminal charge effects

To quantify the terminally weighted charge distribution, we defined the Net Charge Preference at chain Terminus (NCPT). It is calculated as 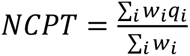, where *q_i_* is the charge of residue *i* in units of *e,* and *w_i_* is a position-dependent weight that assigns higher value to residues near the chain termini. It was chosen as an exponential function 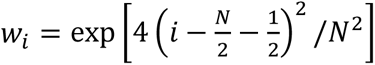 based on the sequence-position-dependent interfacial enrichment measured in charge-free homopolymer simulations (Fig. S1). NCPT therefore represents a terminally weighted measure of sequence charge rather than the total net charge of the chain. Because chain termini are preferentially enriched at the interface, the sign and magnitude of NCPT predict the interfacial charge preference. The NCPT values for sequences A1, A2 and A3 are 0, 0.0089, and -0.0089, respectively, which exhibit a negative correlation with the interphase potential ΔΦ.

To further elucidate the impact of NCPT, we designed two additional sets of sequences based on A2. The first set of sequences includes sequences B1, B2, B3, which contain one, two and three additional neutral beads on the C terminus compared to A2 (Fig. 6A). The addition of these neutral beads results in larger NCPT values 0.0142, 0.0175 and 0.0194 for of B1, B2 and B3. Correspondingly, the interphase potential ΔΦ decreases as NCPT increases, maintaining a robust negative correlation (Fig. 6B C E).

**Figure 6.**
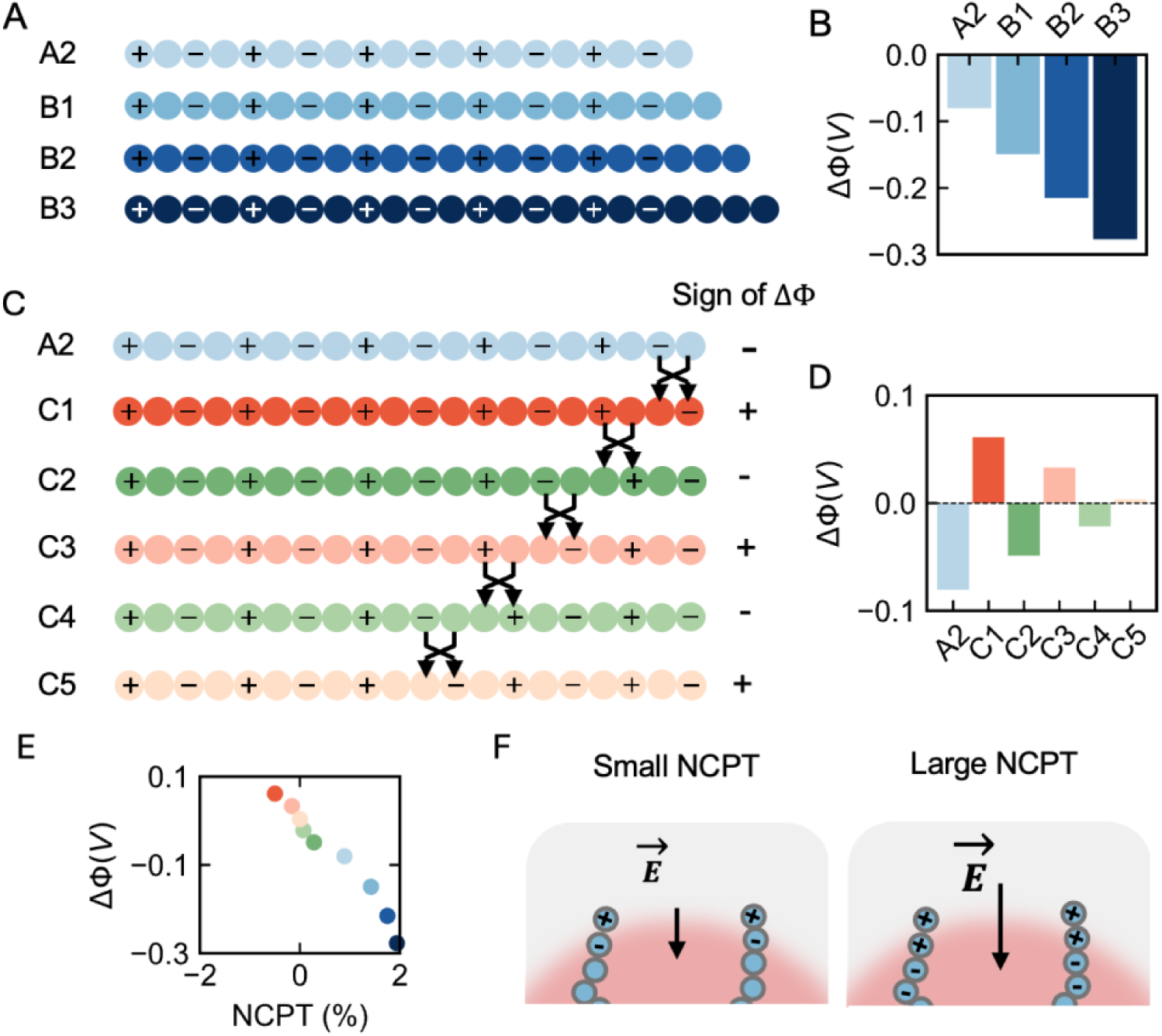
Net Charge Preference at Chain Termini quantitatively captures terminal charge-patterning effects. **(A)** Designed sequence sets used to vary terminal charge patterning by altering the number or placement of neutral beads near the C terminus of A2. These modifications change terminal charge preference while preserving the relevant sequence constraints described in the text. **(B)** ΔΦ values for the B-series sequences. Increasing the terminally weighted positive charge decreases ΔΦ. **(C)** Designed sequence sets used to vary terminal charge patterning by systematically exchanging charged beads with neighboring neutral beads. **(D)** ΔΦ values for the C-series sequences. **(E)** Relationship between ΔΦ and NCPT for the designed sequence sets. The data follow a common negative correlation, demonstrating that terminally weighted charge placement captures both the magnitude and sign of the interphase potential across the sequences examined. **(F)** Schematic showing how terminal charge patterning affects interphase potential.

The second set of sequences was generated through a systematic swapping procedure. First, we swapped terminal charged bead of sequence A2 with its adjacent neutral neighbor to generate sequence C1 (Fig. 6C). We then iteratively swapped the next charged bead with its left neutral neighbor to get sequences C2-C5. Interestingly, the NCPT values alternate its sign and decreases in magnitude with each successive swap. Strikingly, the interphase potential ΔΦ mirrors this behavior, alternating its sign after each swap and its magnitude decreases to zero as the NCPT approaches zero (Fig. 6D). Once again, we observed a negative correlation between NCPT and ΔΦ (Fig. 6E).

Importantly, we noted that the correlation between NCPT and ΔΦ for these two sets of sequences follows a master curve (Fig. 6E). Taken together, these results demonstrate that the NCPT is a crucial determinant of the interphase potential.

### Interplay between interaction asymmetry and NCPT in determining interphase potential

The preceding results establish that the interphase potential is influenced by two distinct mechanisms: 1) asymmetric protein-cation and protein-anion interactions, and 2) terminal charge patterning, as quantified NCPT. We next investigated the interplay between these two factors by simulating the phase separation of A2 sequence with different protein-ion interactions: 1) symmetric interactions (*ε_p_*_+_ = *ε_p_*_−_), 2) stronger protein-cation interactions (*ε_p_*_+_ > *ε_p_*_−_), 3) stronger protein-anion interactions (*ε_p_*_+_ < *ε_p_*_−_) (Fig. 7A).

**Figure 7.**
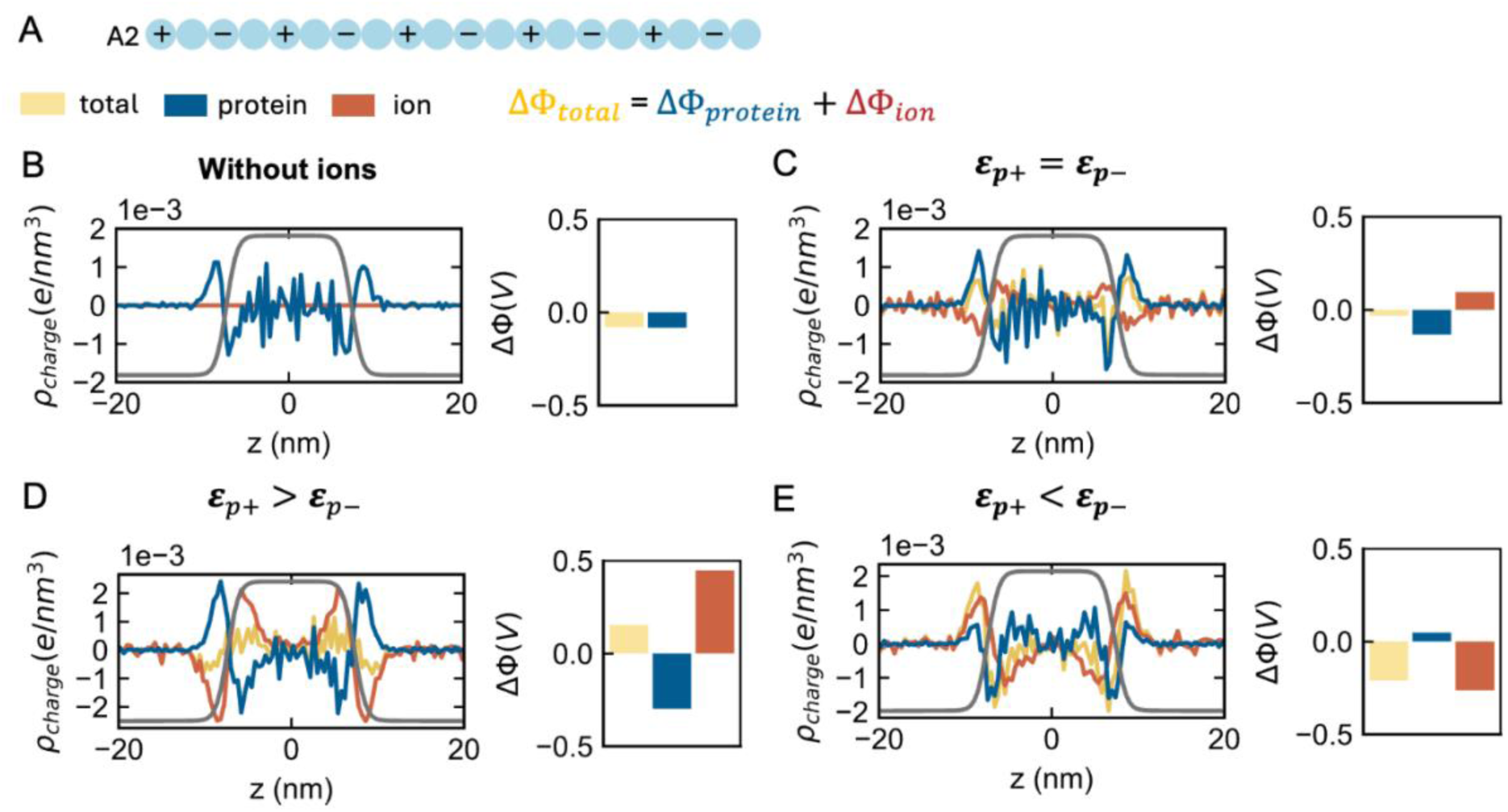
Protein–ion interaction asymmetry competes with terminal charge patterning to determine the interphase potential. **(A)** Schematic of condensates formed by the A2 sequence under three protein–ion interaction conditions: symmetric interactions, *ε_p_*_+_ = *ε_p_*_−_; stronger protein– cation interactions, *ε_p_*_+_ > *ε_p_*_−_; and stronger protein–anion interactions, *ε_p_*_+_ < *ε_p_*_−_. **(B)** In the absence of added ions, terminal charge patterning generates an interfacial protein-charge distribution and a negative ΔΦ. **(C)** With ions that interact symmetrically with the protein, ion redistribution partially compensates for the protein-generated interfacial charge profile, reducing the magnitude of ΔΦ. **(D)** When protein–cation interactions are stronger than protein–anion interactions, preferential cation accumulation opposes the NCPT-generated potential. The ion-mediated contribution dominates under the condition shown, reversing the interfacial charge organization and producing a positive ΔΦ. **(E)** When protein–anion interactions are stronger, preferential anion accumulation reinforces the sequence-generated potential and increases its magnitude. Together, these results show that interaction-encoded and sequence-encoded asymmetries can screen, reinforce, or reverse the interphase potential.

As discussed above, ΔΦ for condensates formed by A2 in the absence of ions is negative, driven solely by the terminal charge patterning of the protein (Fig. 7B). When ions with symmetric interactions are introduced, they tend to neutralize protein-induced interfacial net charge profile. This attenuates the electric double layer (Fig. 7C), and decreases the overall magnitude of ΔΦ, suggesting a screening effect of ions. Conversely, when *ε_p_*_+_ > *ε_p_*_−_, the strong protein-cation interactions cases the direction of the interfacial electric double layer flipped, which leads to a positive ΔΦ (Fig. 7D). In this scenario, the asymmetric protein-ion interaction compete withs the NCPT effect, and ultimately dominates. Similarly, when *ε_p_*_+_ < *ε_p_*_−_, the magnitude of ΔΦ increases compared to the ion-free system, driven primarily by the binding of anions (Fig. 7E). Interestingly, we observed that the net charge profile of proteins has one more peak at the dense-phase side of the interface compared to that formed by ions. This is attributed to the propensity of charge neutrality within the dense phase, and demonstrates the complex interfacial behaviors.

### Sequence net charge modulates the interphase potential

Finally, we examined the influence of overall sequence net charge on ΔΦ using sequence E1, which contains one negative charge at the terminus (Fig. 8A). Counterions were included to maintain overall system neutrality. In addition, we included extra salt ions with asymmetric interactions (*ε_p_*_+_ > *ε_p_*_−_) and varied their concentrations. At low salt concentrations, the negative charges of proteins formed the negatively charged layer on the dense-phase side of the condensate, while the positive counterions formed a compensating positively charged layer on the dilute-phase side (Fig. 8B). This charge separation generates a negative ΔΦ value (Fig. 8C). However, as salt concentration increases, a larger number of cations becomes available for preferential accumulation in the protein-rich phase. The contribution from asymmetric protein–ion interactions therefore becomes progressively more important and eventually reversing the interfacial electric field and leading to a positive ΔΦ value (Fig. 8C). We also varied the position of the negative charge in E1 and observed similar salt-dependent inversion of the interphase potential (Fig. S2), confirming the robustness of this mechanism.

**Figure 8.**
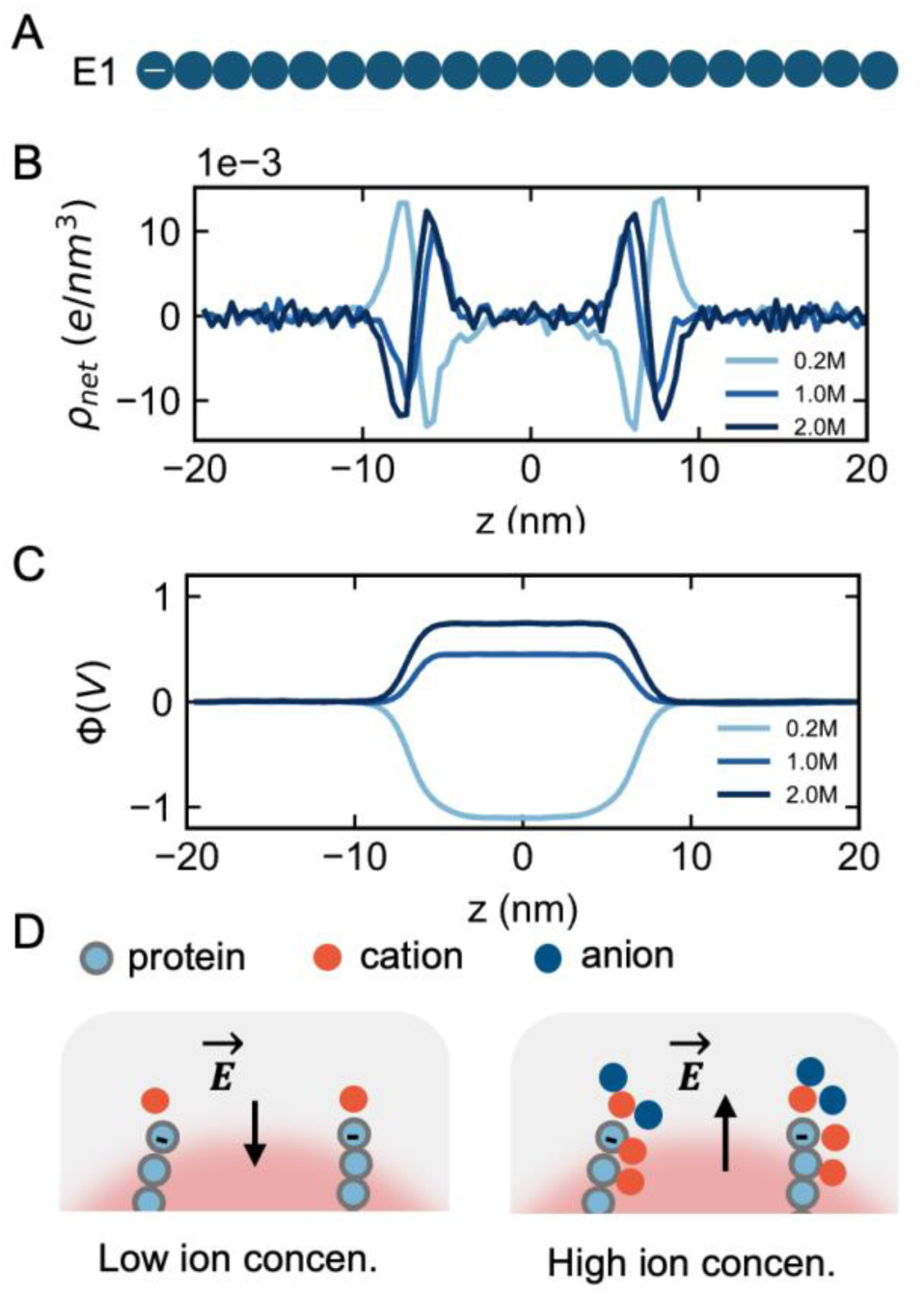
Salt-dependent ion accumulation reverses the interphase potential of a net-charged condensate. **(A)** Schematic of sequence E1, which contains a terminal negative charge and therefore has a nonzero sequence net charge. **(B)** Protein and ion charge-density profiles at representative salt concentrations. At low salt concentration, negatively charged protein sites form a negative layer toward the dense-phase side of the interface, whereas positive counterions form a compensating layer toward the dilute-phase side. At higher salt concentrations, preferential cation accumulation in the protein-rich environment progressively reorganizes and eventually reverses the interfacial charge distribution. **(C)** Interphase electric potential, ΔΦ, as a function of salt concentration. The potential is negative at low salt concentration but increases with salt concentration, passes through zero, and becomes positive when the contribution from asymmetric protein–ion interactions exceeds that generated by the sequence net charge and its counterions. Similar behavior is observed when the negative charge is placed at other sequence positions, demonstrating the robustness of the salt-dependent potential inversion. **(D)** Schematic showing how salt concentration regulates ΔΦ.

## Discussion and Conclusion

In this study, we used a minimalist phenomenological model to identify the physical principles that govern the interphase electric potential difference in model protein condensates. By reducing molecular complexity, this coarse-grained model allowed us to isolate factors that are strongly coupled in real condensates and examine their individual and combined effects on ΔΦ. We focused on two different molecular asymmetries: unequal protein–cation and protein–anion interactions, and sequence-encoded charge patterning near the chain termini. This approach enabled us to determine the minimal conditions under which each mechanism generates interfacial charge separation.

A central finding of this work is that asymmetric protein–ion interactions are sufficient to generate an interfacial electric double layer and a finite ΔΦ in condensates formed by charge-free proteins ^37^. These results complement previous experimental and atomistic studies showing that cations and anions can partition differently between condensate phases and that charge-free protein condensates can nevertheless exhibit interphase potential ^37^. The interaction parameters *ε_p_*_+_ and *ε_p_*_−_, however, should be interpreted as effective affinities rather than as specific interactions. At the molecular level, these effective affinities may incorporate contributions from ion hydration, local coordination, solvent-mediated interactions, and the chemical composition of amino acids in the protein. The model therefore demonstrates the physical consequences of unequal protein-ion affinities but does not resolve their complete chemical origins.

Dense-phase protein volume fraction emerged as an important collective variable controlling the ion-mediated interphase potential. Across variations in chain length, protein–protein interaction strength, and salt concentration, ΔΦ followed a common relationship with the dense-phase protein volume fraction, *φ_p_*. In the moderate-concentration regime, the approximately linear dependence of ΔΦ on *φ_p_* is consistent with previous thermodynamic results showing that ion-transfer free energies vary approximately linearly with protein density. This finding suggests that physically distinct perturbations can regulate condensate electrostatics through their common effects on dense-phase molecular packing and ion-transfer energetics.

At higher protein volume fractions, ΔΦ deviated from the linear relationship with *φ_p_* and eventually decreased despite a continued increase in *φ_p_*. Several effects may contribute to this turnover. Strong protein–protein attraction can reduce the free volume available to ions, promote ion exclusion from the dense phase, or limit ion accessibility to favorable protein-ion interaction sites. High macromolecular density may also alter the interfacial width and reorganize the relative distributions of cations and anions. Alternatively, the ion-transfer free energies may depend nonlinearly on protein concentration in this regime. Further calculations of ion partition coefficients, protein–ion coordination, interfacial charge accumulation, and excess transfer free energies will be needed to distinguish among these mechanisms. Therefore, the common curve observed here should be regarded as an empirical relationship across the conditions examined, rather than a universal equation of state applicable to all condensates.

Chain termini are topologically distinct from interior residues because they have fewer chain-connectivity constraints. This distinction gives terminal residues an enhanced propensity to occupy the interface ^57^. Consequently, charges positioned near the chain termini can contribute more strongly to the interfacial charge-density profile than charges located in the chain interior, thereby modulating both the magnitude and polarity of ΔΦ. Importantly, this mechanism does not require a nonzero sequence net charge. Instead, it arises when positive and negative residues occupy nonequivalent positions relative to the interfacial enriched chain termini, allowing a neutral polyampholyte to generate spatial charge separation. Previous studies have established that the linear patterning of oppositely charged residues regulates IDP conformations, phase behavior, and condensate organization ^62–65^. Existing charge-patterning descriptors, including *κ* ^62^ and “sequence charge decoration” (SCD) ^66^, generally characterize the global arrangement of charges along a sequence, but they do not explicitly account for the enhanced interfacial weighting of residues near the chain termini.

To quantify this terminally weighted charge distribution, we introduced the Net Charge Preference at chain Terminus (NCPT), a terminally weighted sequence descriptor that captures the relative positioning of positive and negative residues near the chain ends. It provides information that is not captured by sequence net charge or terminal residue identity alone. The NCPT weighting function represents the terminal localization preference within the present coarse-grained model, and its quantitative form may depend on chain length, stiffness, interaction strength, interfacial width, or protein architecture. Natural proteins may also contain folded domains, post-translational modifications, or heterogeneous interaction motifs that alter terminal localization. NCPT should therefore be regarded as a physically motivated sequence descriptor validated across the sequence libraries examined here. Its primary purpose is to demonstrate how terminal charge patterning can be translated into a quantitative measure of interfacial electrostatic bias. Because ΔΦ is governed by coupled contributions from sequence patterning, protein–ion interactions, phase composition, and solution conditions, NCPT alone is not expected to predict the interphase potential across all condensate systems. Nevertheless, it provides a useful starting point for developing more general sequence-based models of condensate electrochemical properties.

The competition among terminal charge patterning, protein–ion interaction asymmetry, and overall sequence net charge further demonstrates that condensate electrostatics cannot be inferred from any single molecular property. More generally, combined experimental and coarse-grained studies have shown that sequence patterning and macromolecular architecture can act cooperatively to regulate condensate composition and multiphase organization ^67^. Our results extend this principle to condensate electrostatics by showing that sequence architecture and ion-specific interactions jointly determine ΔΦ. When sequence-encoded and ion-mediated contributions favor the same polarity, they reinforce one another and increase the magnitude of ΔΦ. When they favor opposite directions, they compete, potentially producing cancellation or direction reversal. The salt-dependent inversion observed for the net-charged sequence illustrates this competition: increasing the salt concentration enhances the contribution of preferentially accumulated ions until it overcomes the potential generated by the fixed protein charges and their counterions. Thus, the electrochemical polarity of a condensate emerges from the collective organization of protein charges, mobile ions, and sequence-specific interfacial preferences rather than from protein net charge alone.

The ability to modulate the magnitude and polarity of ΔΦ may have broader consequences for condensate function. Interphase potentials and interfacial electric fields can influence the partitioning of charged metabolites, proteins, nucleic acids, and small molecules. They also modulate local ion composition, pH, redox chemistry, hydrolysis reactions, and interactions between condensates and other cellular structures. Our results further suggest that terminal mutations, alternative splicing, proteolytic processing, or post-translational modifications could alter condensate electrostatics by changing terminal charge patterning, even when the overall sequence net charge changes only weakly. Likewise, changes in salt composition or ion availability could reverse the electrochemical polarity of a condensate without altering its protein sequence. These implications remain experimentally testable predictions rather than direct demonstrations of cellular function.

Several limitations of the present model should be considered. First, the implicit-solvent representation does not explicitly capture water orientation, ion hydration, dielectric heterogeneity, and polarization effect. These effects may contribute to electric potential profiles of real protein condensates. Second, the charge-free homopolymers and designed polyampholytes lack residue-specific interactions, secondary structure, folded domains, and the compositional complexity of multicomponent cellular condensates. Third, our simulations describe equilibrium phase coexistence and do not account for active reactions, sustained ion fluxes, chemical gradients, or condensate aging. Finally, Finally, although explicit ions capture heterogeneous ion distributions and ion-mediated screening, electrostatic interactions are evaluated using a uniform effective dielectric constant because water is represented implicitly. The model therefore does not explicitly resolve differences in dielectric response among the dense phase, dilute phase, and interface ^68, 69^, although some solvent-mediated effects may be incorporated into the effective interaction parameters. Accordingly, the calculated potentials are most appropriately interpreted in terms of their signs, relative trends, and physical mechanisms rather than as quantitatively exact predictions of experimental voltages.

The framework developed here yields several experimentally testable predictions. First, changing ion identity to reverse the relative protein affinities of cations and anions should reverse the direction of ΔΦ in condensates formed by charge-free IDPs. Second, sequences with identical composition and net charge but opposite terminal charge biases should generate potentials of opposite signs. Third, within the moderate-concentration regime, perturbations that alter dense-phase protein volume fraction should produce approximately predictable changes in ΔΦ. Fourth, when sequence-encoded and solution-encoded mechanisms oppose one another, changing salt concentration should produce a condition at which ΔΦ vanishes, followed by direction inversion. These predictions could be tested using systematic sequence variants, salt-series measurements, ion-activity measurements, electrochemical potentiometry, and interface-sensitive spectroscopic methods.

In conclusion, our findings establish molecular asymmetry as a unifying principle underlying electrochemical polarization in model protein condensates. Such asymmetry can be encoded in the relative affinities of proteins for cations and anions or in the placement of charged residues relative to interfacial enriched chain termini. Dense-phase protein concentration controls the collective strength of the ion-mediated mechanism, whereas NCPT provides a physically motivated measure of the terminal charge-patterning contribution. The interplay among these molecular features enables the magnitude and polarity of ΔΦ to be tuned through either sequence design or solution conditions. This framework links molecular interactions and polymer topology to mesoscale electrochemical behavior and provides testable principles for understanding and engineering the electrostatic properties of biomolecular condensates.

## Methods

All slab simulations were performed using OpenMM 8.4.0 package ^70^. The simulation box size was 10*10*40 nm^3^ (if not otherwise specified) with periodic boundary condition. Different number of polymer chains and ions were randomly put into the simulation box and then were subjected to energy minimization. A Langevin integrator with a time step of 10 fs and a friction coefficient of 0.5 ps^-1^ was used to maintain the system temperature at 300K. In order to initialize a slab geometry, we applied a harmonic restrain potential to all the particles along the z-axis for 100 ps, with a force constant of 1 kJ/(mol*nm^2^) and z = 0 as the equilibrium position. After this, we removed the restrain and run for 6 μs. The configurations were saved every 50 ps. The initial 50 ns was discarded as equilibration and the rest was used for subsequent analysis.

The polymer was modeled as a chain of 20 beads (if not otherwise specified) connected by harmonic potentials. The ions were modeled as single beads carrying unit charge. Nonbonded interactions between different pairs of beads contained Lennard-Jones (LJ) potential and Coulomb electrostatic potential. The total energy of the system is written as *E_total_*= *E_bonded_*+ *E_LJ_* + *E_coul_*, and 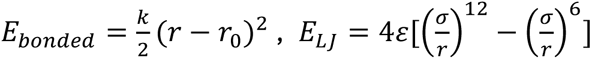, 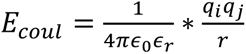, where *r_0_* is 0.381 nm, and *k* is 8033 kJ/(mol*nm^2^). *ε* is the well depth of LJ potential and *σ* is the van der Waals radius. Their values are given in the tables in the Supplementary Information for different system settings. *ε*_0_ is vacuum permittivity, and *ε_r_* is dielectric constant with a value of 80 to model the water screening effect. Particle mesh Ewald (PME) ^71, 72^ was employed to treat the long-range electrostatics with a cutoff distance of 1.5 nm. A scaling factor of 0, 0.5 and 0.5 was applied to 1-2, 1-3 and 1-4 neighbor nonbonded interactions respectively. The molar mass for polymer beads is 57.05 g/mol which mimics that of glycine residue, while the molar mass for ion beads is 22.99 g/mol if not otherwise specified.

## Author Contributions

X.Z. and Y.D. conceived the idea, designed the study, interpreted the simulation results, and wrote the manuscript. F.C. and R.X. performed the simulations and analyzed the results.

## Acknowledgement

This work is supported by the funding from Research Grant Council of Hong Kong (Grant No. 12304126, No. 22302823), Hong Kong Baptist University (RC-FNRA-IG/22-23/SCI/03). All simulations were performed on the MGPU Cluster at the High-Performance Cluster Computing Centre, Hong Kong Baptist University.

