## Supplementary information for "Sequence-dependent molecular asymmetry and architecture define electric potential profiles of biomolecular condensates"

**for**

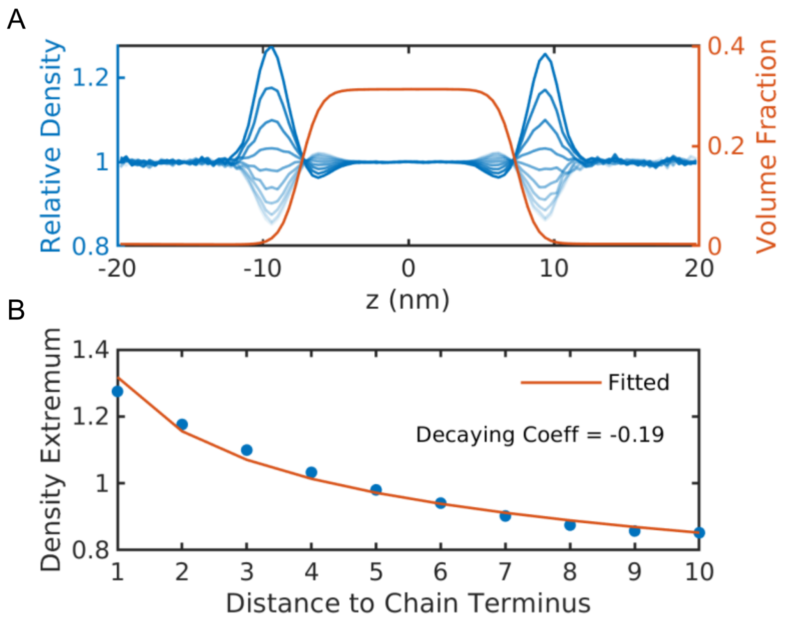


**Figure S1. Charge-free homopolymer model exhibits a sequence-position-dependent interfacial enrichment. (A)** Red curve represents the volume fraction profile of homopolymers. Blue curves from deep to shallow represent the relative density profile of beads at different sequence positions from 1 to 10 counting from chain terminus. Beads closer to chain terminus have a greater propensity at the dilute-phase side of the interface, while beads closer to chain center have a greater propensity at the dense-phase side of the interface. **(B)** Relative density extrema at the dilute-phase side of the interface plotted against the bead's distance to chain terminus. Red curve represents the exponential fitting result.

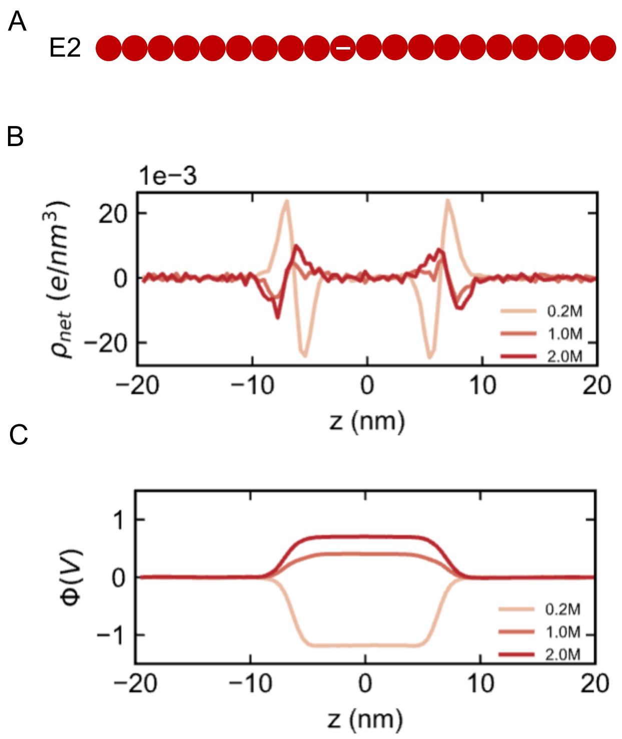


**Figure S2. Salt-dependent ion accumulation reverses the interphase potential of a condensate formed by proteins with non-zero charges. (A)** Schematic of sequence E2, which contains a negative charge at the center and therefore has a nonzero sequence net charge. **(B)** Protein and ion charge-density profiles at representative salt concentrations. At low salt concentration, negatively charged protein sites form a negative layer toward the dense-phase side of the interface, whereas positive counterions form a compensating layer toward the dilute-phase side. At higher salt concentrations, preferential cation accumulation in the protein-rich environment progressively reorganizes and eventually reverses the interfacial charge distribution. **(C)** Interphase electric potential, $\Delta\Phi$, as a function of salt concentration. The potential is negative at low salt concentration but increases with salt concentration, passes through zero, and becomes positive when the contribution from asymmetric protein–ion interactions exceeds that generated by the sequence net charge and its counterions. Similar behavior is observed when the negative charge is placed at other sequence positions, demonstrating the robustness of the salt-dependent potential inversion.

**Table S1. Lennard–Jones (LJ) interaction parameters (ε and σ) used in the coarse-grained molecular dynamics simulations shown in Figure 2 of the main text.** All simulations were performed with 500 polymer chains, 480 cations, and 480 anions. The molar masses of the cation and anion were 22.99 and 35.45 g mol$^{-1}$, respectively.

| $\varepsilon$ | Value (kcal/mol) | $\sigma$ | Value (Å) |
|---|---|---|---|
| $\varepsilon_{++}$ | 0.087 | $\sigma_{++}$ | 2.44 |
| $\varepsilon_{--}$ | 0.036 | $\sigma_{--}$ | 4.48 |
| $\varepsilon_{+-}$ | 0.056 | $\sigma_{+-}$ | 3.46 |
| $\varepsilon_{pp}$ | 0.200 | $\sigma_{pp}$ | 4.50 |
| $\varepsilon_{p+}$ | 0.142 | $\sigma_{p+}$ | 3.47 |
| $\varepsilon_{p-}$ | 0.142 | $\sigma_{p-}$ | 4.49 |

**Table S2. Lennard–Jones (LJ) interaction parameters (ε and σ) used in the coarse-grained molecular dynamics simulations shown in Figure 3 of the main text.** All simulations were performed with 500 polymer chains, 480 cations, and 480 anions.

| $\varepsilon$ | Value (kcal/mol) | $\sigma$ | Value (Å) |
|---|---|---|---|
| $\varepsilon_{++}$ | 0.087 | $\sigma_{++}$ | 2.44 |
| $\varepsilon_{--}$ | 0.087 | $\sigma_{--}$ | 2.44 |
| $\varepsilon_{+-}$ | 0.087 | $\sigma_{+-}$ | 2.44 |
| $\varepsilon_{pp}$ | 0.200 | $\sigma_{pp}$ | 4.50 |
| $\varepsilon_{p+}$ | 0.142 | $\sigma_{p+}$ | 3.47 |
| $\varepsilon_{p-}$ | 0.100, 0.142, 0.180 | $\sigma_{p-}$ | 3.47 |

**Table S3. Simulation settings for the coarse-grained molecular dynamics simulations shown in Figure 4 of the main text. (A)** Lennard–Jones (LJ) interaction parameters (ε and σ) used in the simulations shown in Figure 4A. **(B)** Polymer chain length, numbers of polymer chains and ions, and simulation box dimensions used in the simulations in Figure 4A. **(C)** LJ interaction parameters (ε and σ) used in the simulations in Figure 4B. All simulations were performed with 500 polymer chains, 480 cations, and 480 anions. **(D)** LJ interaction parameters (ε and σ) used in the simulations in Figure 4C. The number of polymer chains was fixed at 500, whereas the numbers of cations and anions were varied from 48 to 4800 (48, 96, 240, 480, 960, 1920, 2880, 3840, and 4800 for each ion species).

A

| $\varepsilon$ | Value (kcal/mol) | $\sigma$ | Value (Å) |
|---|---|---|---|
| $\varepsilon_{++}$ | 0.087 | $\sigma_{++}$ | 2.44 |
| $\varepsilon_{--}$ | 0.087 | $\sigma_{--}$ | 2.44 |
| $\varepsilon_{+-}$ | 0.087 | $\sigma_{+-}$ | 2.44 |
| $\varepsilon_{pp}$ | 0.200 | $\sigma_{pp}$ | 4.50 |
| $\varepsilon_{p+}$ | 0.142 | $\sigma_{p+}$ | 3.47 |
| $\varepsilon_{p-}$ | 0.100 | $\sigma_{p-}$ | 3.47 |

B

| Chain length | $N_p$ | $N_{+}$ | $N_{+}$ | Box size ($nm^3$) |
|---|---|---|---|---|
| 15 | 667 | 480 | 480 | 10*10*40 |
| 20 | 500 | 480 | 480 | 10*10*40 |
| 30 | 1125 | 1620 | 1620 | 15*15*60 |
| 40 | 844 | 1620 | 1620 | 15*15*60 |
| 60 | 1333 | 3840 | 3840 | 20*20*80 |
| 80 | 1000 | 3840 | 3840 | 20*20*80 |

C

| $\varepsilon$ | Value (kcal/mol) | $\sigma$ | Value (Å) |
|---|---|---|---|
| $\varepsilon_{++}$ | 0.087 | $\sigma_{++}$ | 2.44 |
| $\varepsilon_{--}$ | 0.087 | $\sigma_{--}$ | 2.44 |
| $\varepsilon_{+-}$ | 0.087 | $\sigma_{+-}$ | 2.44 |
| $\varepsilon_{pp}$ | 0.18, 0.20, 0.22, 0.24, 0.26, 0.28, 0.30, 0.40, 0.60, 0.80, 1.00 | $\sigma_{pp}$ | 4.50 |

| | | | |
|---|---|---|---|
| $\varepsilon_{p+}$ | 0.142 | $\sigma_{p+}$ | 3.47 |
| $\varepsilon_{p-}$ | 0.100 | $\sigma_{p-}$ | 3.47 |

D

| $\boldsymbol{\varepsilon}$ | **Value (kcal/mol)** | $\boldsymbol{\sigma}$ | **Value (Å)** |
|---|---|---|---|
| $\varepsilon_{++}$ | 0.087 | $\sigma_{++}$ | 2.44 |
| $\varepsilon_{--}$ | 0.087 | $\sigma_{--}$ | 2.44 |
| $\varepsilon_{+-}$ | 0.087 | $\sigma_{+-}$ | 2.44 |
| $\varepsilon_{pp}$ | 0.200 | $\sigma_{pp}$ | 4.50 |
| $\varepsilon_{p+}$ | 0.142 | $\sigma_{p+}$ | 3.47 |
| $\varepsilon_{p-}$ | 0.100 | $\sigma_{p-}$ | 3.47 |

**Table S4. Lennard–Jones (LJ) interaction parameters (ε and σ) used in the coarse-grained molecular dynamics simulations shown in Figures 5, 6, and 7B of the main text**. All simulations were performed with 500 polymer chains and no ions.

| $\boldsymbol{\varepsilon}$ | **Value (kcal/mol)** | $\boldsymbol{\sigma}$ | **Value (Å)** |
|---|---|---|---|
| $\varepsilon_{pp}$ | 0.200 | $\sigma_{pp}$ | 4.50 |

**Table S5. Lennard–Jones (LJ) interaction parameters (ε and σ) used in the coarse-grained molecular dynamics simulations shown in Figure 7C–E of the main text.**

The three $(\varepsilon_{p+}, \varepsilon_{p-})$ parameter sets correspond to Figure 7C, 7D, and 7E, respectively. All simulations contained 500 polymer chains, 480 cations, and 480 anions.

| $\boldsymbol{\varepsilon}$ | Value (kcal/mol) | $\boldsymbol{\sigma}$ | Value (Å) |
| --- | --- | --- | --- |
| $\varepsilon_{++}$ | 0.087 | $\sigma_{++}$ | 2.44 |
| $\varepsilon_{--}$ | 0.087 | $\sigma_{--}$ | 2.44 |
| $\varepsilon_{+-}$ | 0.087 | $\sigma_{+-}$ | 2.44 |
| $\varepsilon_{pp}$ | 0.200 | $\sigma_{pp}$ | 4.50 |
| $\varepsilon_{p+}, \varepsilon_{p-}$ | 0.142, 0.142<br>0.142, 0.100<br>0.100, 0.142 | $\sigma_{p+}$<br>$\sigma_{p-}$ | 3.47<br>3.47 |

**Table S6. Lennard–Jones (LJ) interaction parameters (ε and σ) used in the coarse-grained molecular dynamics simulations shown in Figure 8 of the main text.** The number of polymer chains was fixed at 500. The numbers of cations and anions were set to 480, 2400, and 4800 for each ion species.

| $\varepsilon$ | Value (kcal/mol) | $\sigma$ | Value (Å) |
|---|---|---|---|
| $\varepsilon_{++}$ | 0.087 | $\sigma_{++}$ | 2.44 |
| $\varepsilon_{--}$ | 0.087 | $\sigma_{--}$ | 2.44 |
| $\varepsilon_{+-}$ | 0.087 | $\sigma_{+-}$ | 2.44 |
| $\varepsilon_{pp}$ | 0.240 | $\sigma_{pp}$ | 4.50 |
| $\varepsilon_{p+}$ | 0.300 | $\sigma_{p+}$ | 3.47 |
| $\varepsilon_{p-}$ | 0.100 | $\sigma_{p-}$ | 3.47 |
